# Light-Guided Rabies Virus Tracing for Neural Circuit Analysis

**DOI:** 10.1101/2023.03.04.531104

**Authors:** Shuyang Zhang, Yunhan Ma, Wasu Ngamkanjanarat, Sola Takahashi, Daniel Gibbs, Todd Coleman, Sapphire Doan, Phillip Kyriakakis

**Author notes:** Corresponding Author: Phillip Kyriakakis.

## Abstract

Neuronal tracing methods are essential tools to understand the fundamental architecture of neural circuits and their connection to the overall functional behavior of the brain. Viral vectors used to map these transsynaptic connections are capable of cell-type-specific and directional-specific labeling of the neuronal connections. Herein, we describe a novel approach to guide the transsynaptic spreading of the Rabies Virus (RV) retrograde tracer using light. We built a Baculovirus (BV) as a helper virus to deliver all the functional components necessary and sufficient for a nontoxic RV to spread from neuron to neuron, with a light-actuated gene switch to control the RV polymerase, the *L gene*. This design should allow for precisely controlled polysynaptic viral tracing with minimal viral toxicity. To use this system in a highly scalable and automated manner, we built optoelectronics for controlling this system *in vitro* with a large field of view using an off-the-shelf CMOS sensor, OLED display panel, and microcontrollers. We describe the assembly of these genetic circuits using the uLoop DNA assembly method and a library of genetic parts designed for the uLoop system. Combining these tools provides a framework for increasing the capabilities of nontoxic tracing through multiple synapses and increasing the throughput of neural tracing using viruses.

## INTRODUCTION

In the nervous system, information transfer or signal propagation occurs through a complex of neural circuits composed of 70 million neurons in the mouse brain (1) and 86 billion neurons (2) in the human brain that makes around 10^15^ neural connections (3). These connections form specific neural pathways to achieve essential cognitive functions such as information processing and memory storage (4). To understand the relationship between neural circuits and their corresponding functions, their connections can be observed by implementing neural tracing methods. By identifying specific neural circuits, researchers can build empirical models of the nervous system and study the pathological cause of many psychiatric diseases (5). As a result, the development and optimization of many kinds of neural tracing methods have been ongoing since the 1970s (6).

Kristensen and Olsson introduced the central principle that most modern neural tracers use in 1971 (7,8). They recognized that the molecular transporting mechanism along the axons of the nerve cells can also carry chemical substances with cell-labeling potentials (8). This intrinsic cellular cargo movement became the mainstream approach for tracers to travel within neurons. Traditionally isotope-labeled amino acids were used as tracing units (9,10), however, these tracers relied on bulky autoradiographic instruments for detection (10). Now, most neuron tracing methods use light and electron microscopy to image the tracers. Older microscopic tracing methods used dyes, such as Evans Blue dye (EB) or fluorescently tagged beads (11) that are injected directly into neural sites, but these are diluted during transport and degraded over time (12). In contrast, viral tracers such as Herpes Simplex Virus (HSV), Adeno-Associated Virus (AAV), and Rabies Virus (RV) efficiently enter neurons and travel anterogradely, retrogradely, or both. Unlike older methods, these viral tools have can express reporter proteins that can be imaged with light and electron microscopy with genetic/cell type specificity. In addition, they can express tools such as opsins or Designer Receptors Exclusively Activated by Designer Drugs (DREADDS) that can modulate the neural activity of the labeled cells (13,14). Depending on which viral tracer is used, the neuron labeling can be (1) static, only labeling neurons with initial contact, (2) monosynaptic, labeling the neurons that have a direct synaptic connection with the initial neuron, and (3) polysynaptic, labeling multiple orders of synaptically connected neurons. The combination of viral tracers with transgene expression systems, such as Cre-LoxP, provides an added layer of control over where the tracers spread, allowing for tracing single synaptic connections (15). Specific properties, such as genetic payload size, cellular transport, toxicity, and transgene expression levels, make different viral tracers suitable for different purposes (9). For example, AAV can be packaged with around 5kb of transgenes (16), lentivirus can be packed with approximately 8kb (16), and with the rabies glycoprotein and *L* gene removed RV can package up to 7.5kb if rescued (17,18). Among the vectors with the largest transgene capacity that infect neurons are Herpes Simplex Virus (HSV) and baculovirus (BV), which allow 50kb or more transgenes (19). Therefore, HSV and baculovirus are the most ideal for using genetic tools that require large genes or genetic circuits. However, HSV and BV have limitations of high cellular toxicity or inability to be directly used as a tracer, respectively.

Recombinant RV is a popular tool for neural tracing largely because the spread of the virus can be controlled by deletion of the gene necessary for spreading, the rabies virus glycoprotein (RABV-G). Importantly, the deletion of RABV-G can be rescued by expressing it *in trans* to enable the controlled trans-synaptic spread of RV (20). This powerful approach allows for monosynaptic tracing, where the virus only spreads to neurons directly synaptically connected to the primary RV-infected neuron. The avian viral receptor (TVA) that facilitates RV transsynaptic spread provides another level of genetic control. Without these controls the virus would spread continuously, infecting the entire neural network and making it difficult to determine what connections are direct or indirect. To achieve specificity of monosynaptic tracing of RVΔG strains, helper viruses (such as AAV or Lentivirus) or transgenic animals can complement RABV-G and/or TVA, into desired neuron or neuron types (20).

Soumya Chatterjee and others introduced mutant RV vectors with the deletion of both *G* and *L* genes (RVΔGL) to minimize cellular toxicity and allow for long-term neural labeling (18). Compared to the traditional RVΔG tracing vector, this variant tuned down the viral genome replication rate by additionally knocking out the viral polymerase, the *L* gene (21). This advance showed that the RV strain was nontoxic and could infect and label neurons effectively. However, the large size of the *L* gene made it challenging to rescue RV’s ability to spread because the glycoprotein (G, or RABV-G), TVA, and the L protein all must be complemented *in trans* (18). To construct the complement system for controlling the spreading of this tracer, BV with its large payload capacity has the potential to complement these genes as well as additional tools, such as a fluorescent Cre reporter and optogenetic tools.

Many photoreceptor proteins have been used as optogenetic tools to control neural activity or other cellular activities, such as gene expression (22,23). By incorporating optogenetics tools in the system, light can guide the timing and location of viral activity (24). Phytochrome photoreceptors possess useful properties that can allow for tight spatio-temporal control of viral activity. For example, the *Arabidopsis thaliana* phytochrome PhyB can be activated by light of one wavelength (650nm red light) and inactivated by light of a different wavelength (750nm Far-Red light) (**Figure 1**) (25–27). Because of this property, it is possible to stimulate a small area with red light by using far-red light to tightly restrict the stimulation to a specific area, even on a subcellular level (28,29).

**Figure 1:**
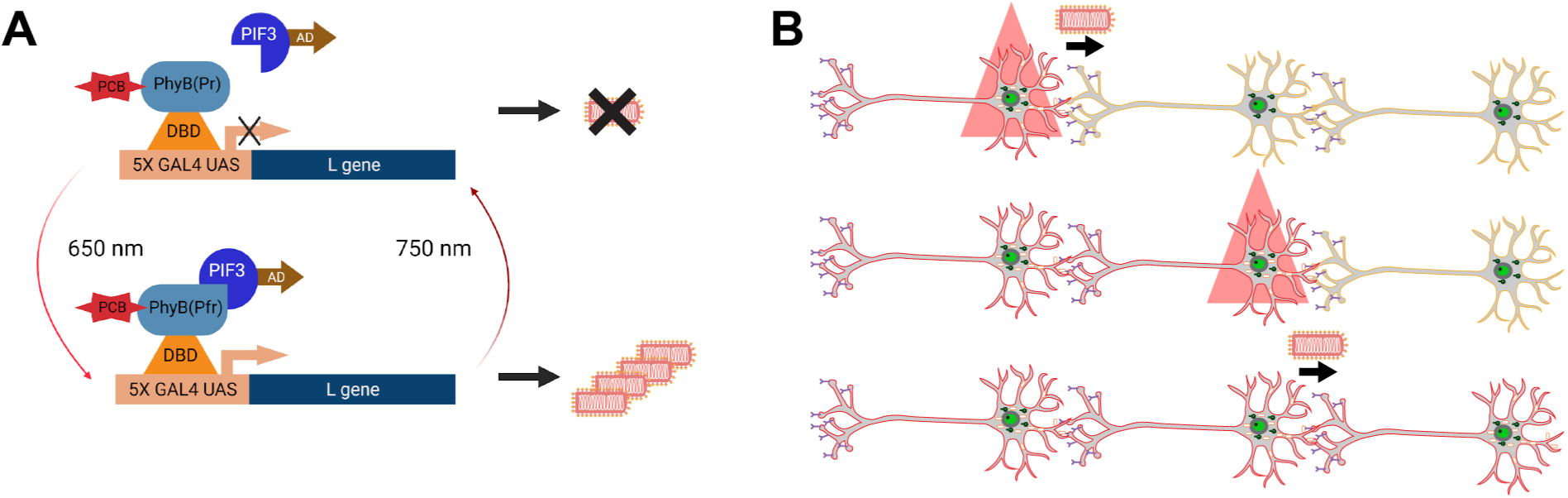
Optically controlled neural tracing. **(A)** The PhyB optogenetic switch controls the expression of the RV *L gene* in a red/far-red light dependent manner. The PhyB photoreceptor is bound to the Gal4 UAS promoter through the DNA-Binding Domain (DBD). The light-dependent interacting protein PIF3 is linked with gene Activation Domain (AD). In the dark, PhyB does not dimerize with PIF3-AD, leaving the *L gene* in the uninduced state. When stimulated by red light (650nm), PhyB changes to Pfr conformation and recruits PIF3-AD to the promoter, activating the *L gene.* This process can be reversed or suppressed with far-red light (750 nm). **(B)** With the *L gene* induced, the RV genome is replicated and RV spreads to synaptically connected neurons, labeling it with mScarlet-CAAX. Through a pulse of red light, the PhyB/PIF3 gene switch will induce *L gene* expression in a second neuron further spreading RV to synapti­cally connected neurons.

PhyB was first shown to control gene activation using a two-hybrid assay with PhyB fused to a DNA-binding Domain (PhyB-DBD) and Phytochrome Interacting Factor 3 (PIF3) fused to a transActivation Domain (PIF3-AD) (**Figure 1**) (**26**). Upon stimulation with red light, a transActivation Domain is recruited to a promoter region, turning a gene on. When stimulated with far-red light (or when incubated in the dark), PhyB-DBD does not interact with PIF3-AD, and the transActivation Domain is not recruited to the promoter, turning a gene off. Since this system can control an arbitrary gene, it can be combined with RV neural tracers to provide genetic control of transsynaptic spreading.

To control PhyB optical stimulation it is possible to use an LED array and combine it with a lensless imaging system to image and control tracing larger areas than a microscope and at a fraction of the cost. Lensless imaging is an emerging technique in microscopy. Unlike conventional fluorescent microscopes, lensless-based imaging systems collect light without an objective, allowing light collection from a larger sample area, essentially the size of the optical sensor. This light pattern is recorded and then processed by pre-calibrated computational algorithms to reconstruct the original image (30). These lensless imaging techniques can acquire images with a significantly larger field-of-view (FOV) compared to conventional microscopes while maintaining a reasonable spatial resolution (31). This technology is well suited for the closed-loop imaging and stimulation of RV tracing because the larger FOV makes it capable of continuously imaging large areas of an entire mouse or rat brain slice and the entire device could be as small as a few cubic centimeters (32,33). Thus, this closed-loop system could image and optically control polysynaptic tracing at a much lower cost and a much lower footprint, compared to using conventional microscopy. In addition, since they can stay inside an incubator while tracing, they would require little hands-on time and could enable automation of mapping specific polysynaptic connections.

In this work, we produced helper viral vectors based on high-capacity BV and integrated the genetic circuitry to produce all the complementary RV proteins with a PhyB/PIF3 optogenetic tool for optical control of the L-gene (25). To implement the use of optogenetics to control viral spreading and trace neural connections, we additionally designed an optoelectronic platform for simulating gene expression for optically controlled polysynaptic tracing. Through miniaturization and automation, this approach to neuronal tracing could enable the tracing of brain slices at scale. Because the system is capable of controlling polysynaptic tracing neural circuits, this approach can pave the way to map complex neural circuits in different developmental contexts or using genetic models of neurological diseases.

## RESULTS

### Optogenetic Baculovirus Helper Virus Design and Construction

We designed the baculovirus to deliver four main functional components: (1) the PhyB/PIF3 optogenetic switch for light controllable gene expression, (2) the *L* gene under the control of the Gal4 UAS promoter, (3) the TVA receptor+RABV-G (TVA+G) for selective neuronal RV entry and spread, and (4) a Cre-dependent fluorescent reporter. The size of this gene combination (PhyB/PIF3 gene switch (6.7kb), the *L* gene (6.7kb), TVA+*G* gene (3.4kb), the Cre dependent reporter (4.1kb), and other components (totaling 26.5kb) far exceeds the transgene limit of commonly used viral vectors (AAV virus (5kb) and lentivirus (8kb)). Since the BV has a 50kb+ genetic payload capacity and has low cytotoxicity in mammalian cells, even at a high Multiplicity of Infection (MOI) (19,34,35), BV makes an ideal high-capacity helper virus system for the nontoxic RVΔGL.

The PhyB optogenetic system regulates the expression of the *L* gene (*i.e.* RV spreading) using red and far-red light (**Figure 1**). To control the *L* gene with light, the PhyB photoreceptor is linked with Gal4 DNA-Binding Domain (Gal4 DBD), which interacts with the PIF3 protein linked with the Activation Domain (AD) in a red/far-red dependent manner. The BV vector contains a Gal4 Upstream Activation Sequence (UAS) and promoter for modulating the *L* gene. Thus, when the PhyB is inactive, it is bound to the promoter, but not to PIF3-AD. Since the AD is not recruited to the promoter, *L* gene expression is off and RVΔGL-Cre will not replicate or spread. Once PhyB is activated by red light, PhyB forms a complex with PIF3-AD, recruiting the AD to the promoter, and inducing *L* gene expression. With the *L* gene on, the RV genes are expressed, allowing it to replicate and spread to synaptically connected neurons.

To restrict the activity of both the BV and RV to neurons, the TVA+G cassette, the PhyB optogenetic tool, and the Cre-dependent fluorescent reporter are all expressed using neural-specific promoters derived from the Human *synapsin 1* gene (36). With this arrangement, only neurons infected with BV will contain the TVA receptor, the fluorescent reporters, and the optogenetic tools for regulating RV spread (**Figure 2**). Once a BV-infected neuron expressing Nuclear Localized mNeonGreen, (NLS-GFP) is subsequently infected with RVΔGL-Cre, Cre will recombine the fluorescent reporter, leading to the expression of membrane-bound mScarlet (mScarlet-CAAX), labeling the entire neuron to highlight pathways within the neural circuit.

**Figure 2:**
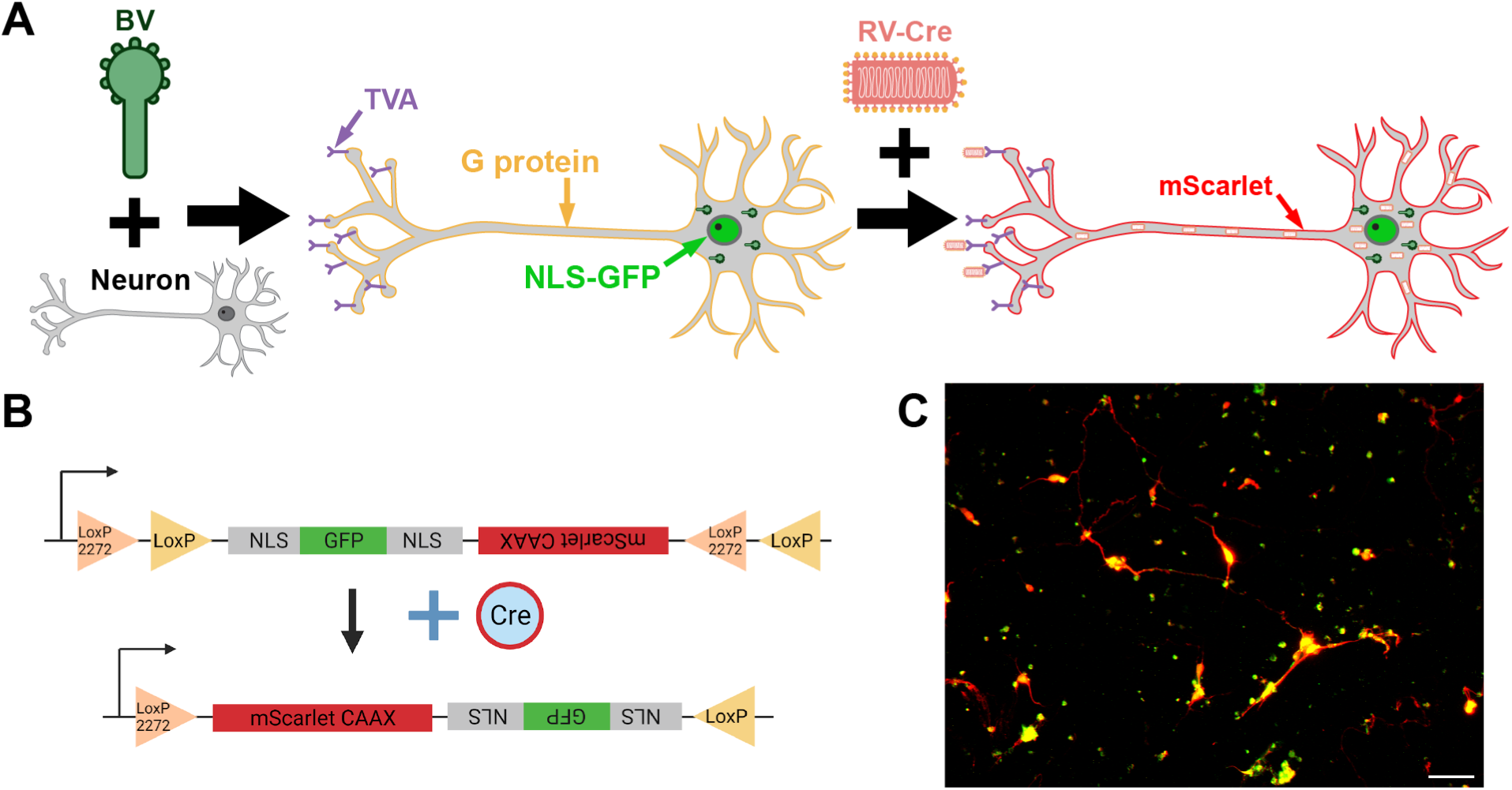
Control and visualization of RVΔGL-Cre neural tracing using a BV helper virus. **(A)** Upon BV infection, TVA+G proteins and NLS-GFP are expressed. BV infected neurons (which contain TVA) further infected with RVΔGL-Cre. Cre recombines the fluorescent Cre reporter, switching from NLS-GFP production to mScarlet-CAAX production. **(B)** The Cre reporter is flanked by DIO LoxP sites. Initially NLS-GFP gene is expressed. After Cre recombi­nation, the DNAorientation is inverted and the mScarlet-CAAX reporter is expressed, highlighting neurons infected with ΔGL-Cre RV. **(C)** Rat E18 cortical neurons electroporated with the Cre reporter and Cre. Neurons with only the Cre reporter show a GFP-NLS signal. Neurons co-transfected with the reporter and Cre are labeled with mScarlet-CAAX. Scale bar = 50um.

For *in vitro* and brain slice experiments, the baculovirus will first be used to infect neurons at a high MOI to infect as many neurons as possible, producing TVA/G proteins, the NLS-GFP reporter, and the PhyB/PIF3 optogenetic switch (**Figure 2A**). After, RVΔGL-Cre is introduced into the neuron samples and can only infect neurons previously infected with the BV (**Figure 2A**). Once inside the neuron, Cre from the RVΔGL-Cre particle recombines with the Cre reporter switching the NLS-GFP expression to mScarlet-CAAX. Since the RVΔGL-Cre is replication-deficient in the absence of *L* gene expression, the viral load will remain nontoxic and unable to infect other neurons. Then, using red light illumination, the expression of the *L gene* is induced through the PhyB/PIF3 optogenetic switch. By complementing the *L* gene with light, RVΔGL-Cre replicates and spreads through retrograde synaptic connections. This process can be repeated to label the next-order connections and can be parallelized to map multiple circuits at once. In addition, by regulating the *L* gene with light, toxic *L* gene expression can be minimized and transient, enabling mapping polysynaptic connections of neural circuits, while constraining the toxicity of the system to single cells.

### Details of the Expression Constructs

The Bac-to-Bac (Bacteria to Baculovirus) system allows the specific DNA to transfer from a bacterial donor plasmid into the baculovirus receiver vector, allowing efficient insertion of large constructs into the BV genome (37). This technique works by transposition of a target DNA/gene flanked with Tn7 transposon elements sequences, Tn7L, and Tn7R on a bacterial plasmid into the attnTn7 docking sites on the baculovirus shuttle vector in bacteria (also known as a bacmid, essentially a very large plasmid or Bacterial Artificial Chromosome (BAC)) (37). This allows for the simple insertion of genes into the BV genome and allows for easy clonal selection before virus production. This work produced three large multi-part plasmids for optically guided tracing designed for easy insertion into the BV genome. The first design includes an optical gene switch based on PhyB that controls the expression of the RV L gene. **(described in detail above and in Figure 3)**. We assembled all of these parts from a library of uLoop parts we constructed **(Figure S2-S6)**. This library includes many individual parts for building genetic circuits including mammalian and insect promoters, insulators [D4Z4c (38), A2 (39–41), Hs4 (41) (42) (43) (44)], fluorescent proteins, and luciferases [mNeonGreen-NLS (45), mScarlet-CAAX (46), EPIC Luciferase (47), Renilla Luciferase (48)], optogenetic parts (25), selection markers (CamR, GentR), polyA tails (SV40 PA, BGH PA, Rabb-beta globin PA), recombination sites (FRT (49), TN7 (50)), new plasmids for the uLoop system and more (**see Tables 1-4**). In addition to the individual Level 0 and Level 1 parts, we included our assembled Level 2, Level 3, and Level 4 parts. We also include a spreadsheet for planning and keeping track of the uLoop assemblies **(Figure S5-S6 and File “uLoop library and planning template.xlsx” in our GitHub (see below))**.

**Figure 3:**
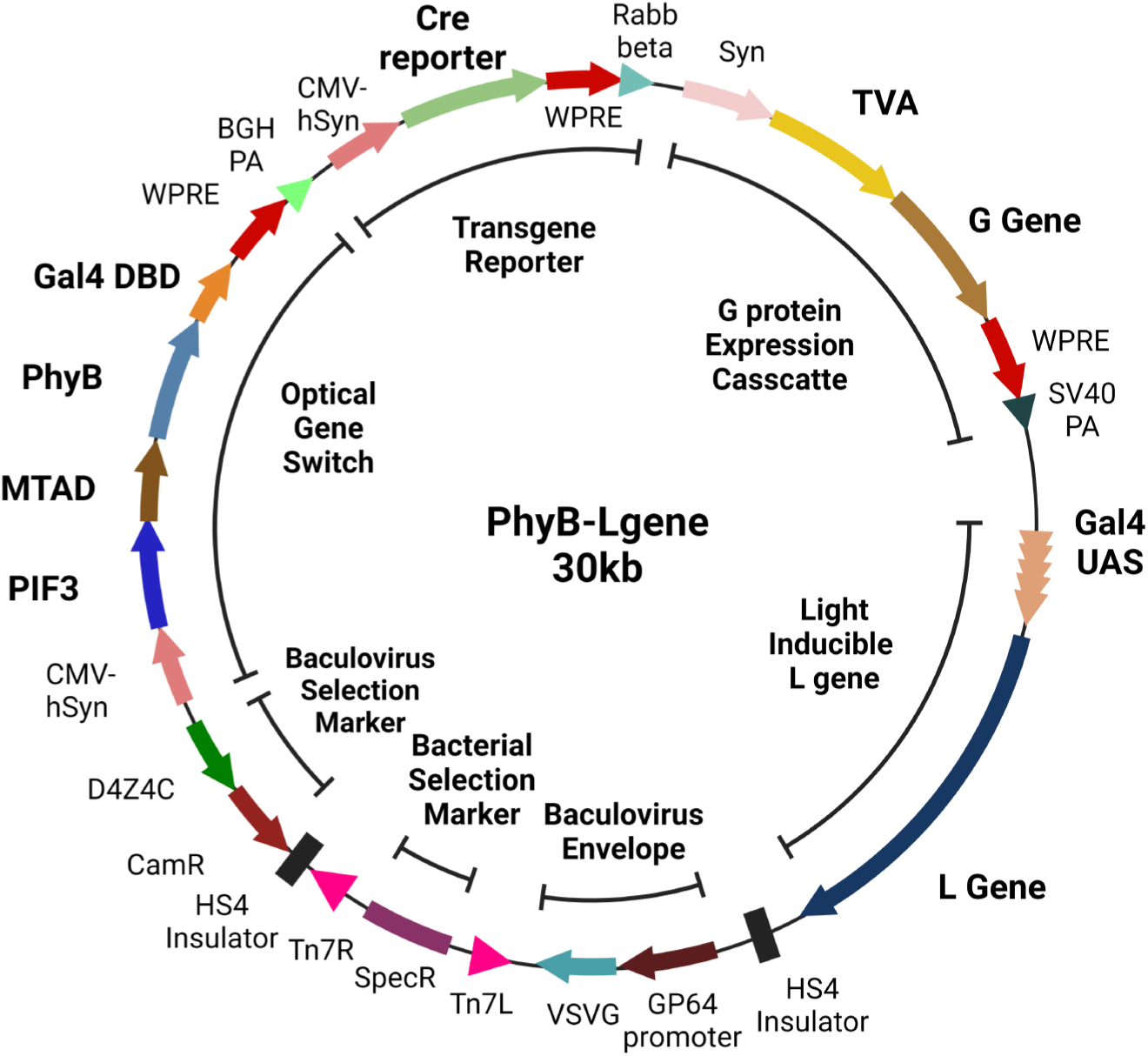
Plasmid map for construct used to insert into the baculovirus genome. The fully assembled PhyB-Lgene plasmid contained seven main functional groups: **(1)** the PhyB optical gene switch, **(2)** the fluorescent Cre reporter, **(3)** the TVA/G protein expression cassette, **(4)** the light-inducible *L gene,* **(5)** a bacterial selection marker for inserting the genes into the BV genome, **(6)** a bacterial selection marker for plasmid cloning, and **(7)** the Vesicular stomatitis virus glycoprotein (VSVG) for enhanced transduction into mammalian cells. The entire baculovirus insertion cassette is flanked by Tn7 transposon sites forTn7-mediated trans­position into the bacmid vector. The size of the resulting plasmid is 29.4kb. (CamR = Chloramphenicol Resistance, D4Z4C = insulator, CMV-hSyn = CMV promoter-en­hancer-human synapsin chimeric promoter, WPRE = Woodchuck hepatitis virus Post-transcrip­tional Response Element, BGH PA = Bovine Growth Hormone PolyA tail, Syn = Synapsin promoter, Rabb beta = Rabbit beta-globin PolyA tail, SV40 PA= Simian Vacuolating virus 40 PolyA tail, SpecR = Spectinomycin Resistance, KanR = Kanamycin Resistance)

### Design of the Stimulation and Imaging System

Similar to the shadow imaging approach, the lensless imaging system developed in this work can be described as two complexes. The core components of these two complexes are a DSLR CMOS sensor for image capture and an Organic Light-Emitting Diode (OLED) light panel for illumination. As shown in the schematic diagram in **Figure 4**, the overall architecture starts with the illumination panel (OLED display) at the bottom of the device, a specimen chamber in the middle, and an imaging sensor at the top. The main purpose of this type of inverted setup compared to common shadow lensless imaging setups is to be able to place the sample as close to the OLEDs as possible to achieve precise activation of the optogenetics system. The specimen is placed directly on top of the OLED with each OLED pixel regulating output intensities and wavelength of the light at a specific region.

**Figure 4:**
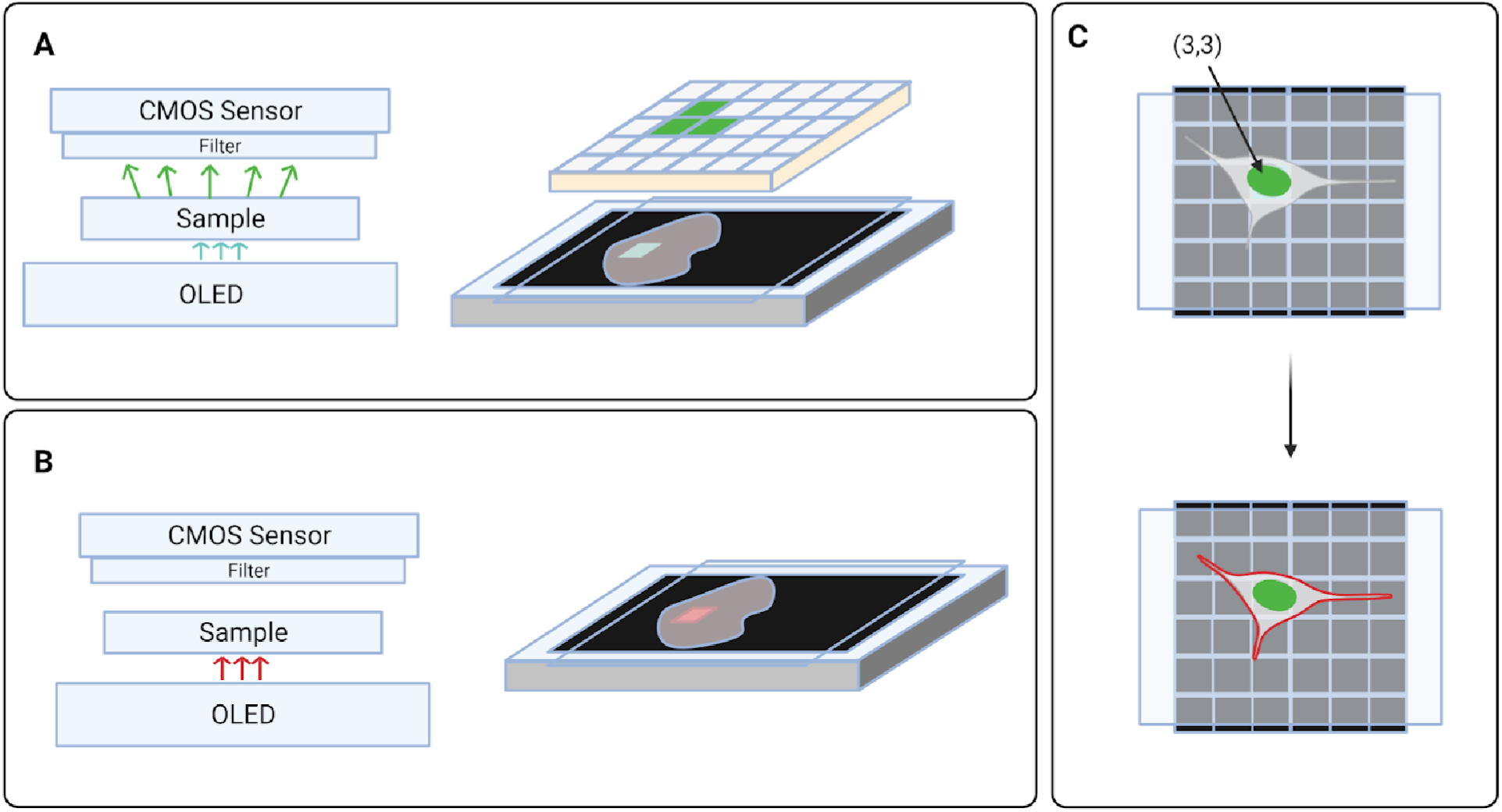
Lensless imaging and optogenetic stimulation system using a CMOS sensor and OLED panel. **(A)** The system architecture represents the horizontal relationship between the OLED panel, sample of interest, emission filter, and the CMOS sensor. Neurons initially infected with BV will produce nuclear GFP signals which can be detected by this setup. **(B)** Red pixels from the OLED panel will activate the PhyB photoreceptor inducing L gene expression and RV spreading to neurons connected to the stimulated cell. **(C)** This setup viewed from the top shows how the single nuclei of a neuron can be stimulated by an OLED (pixel) to activate viral spreading and Cre recombination to label infected cells with high spatial accuracy.

LEDs/OLEDs were chosen for the illumination system because, unlike Liquid Crystal Displays (LCD), it does not require a backlight, minimizing heat and unwanted light leakage. Since the PhyB photoreceptor is highly sensitive to red light activation (saturating at 40nW/cm^2^) (25). For imaging, CMOS sensors are commonly used in lensless imaging. Compared to their counterpart, CCD sensors, CMOS sensors are better suited for lab-on-chip applications because of their faster digital readout speed and lower power consumption rate. In addition, using a CMOS chip extracted from a DSLR camera is widely available and inexpensive. Using this design, sections of the rat’s brain (20.7mm x 15.5 mm) or the mouse’s brain (13.7mm x 5.5mm) can be used. The large active pixel area (∼ 20mm x 20mm) of a standard DSLR CMOS sensor provides sufficient effective sensing to capture the entire cross-sectional area in one frame. However, several sensors could be easily combined and illumined using a larger LED panel (*e.g.* an iPad) for imaging and tracing in larger brains. The overall configuration of the integrated illumination and image acquisition is shown in **Figure 5**.

**Figure 5:**
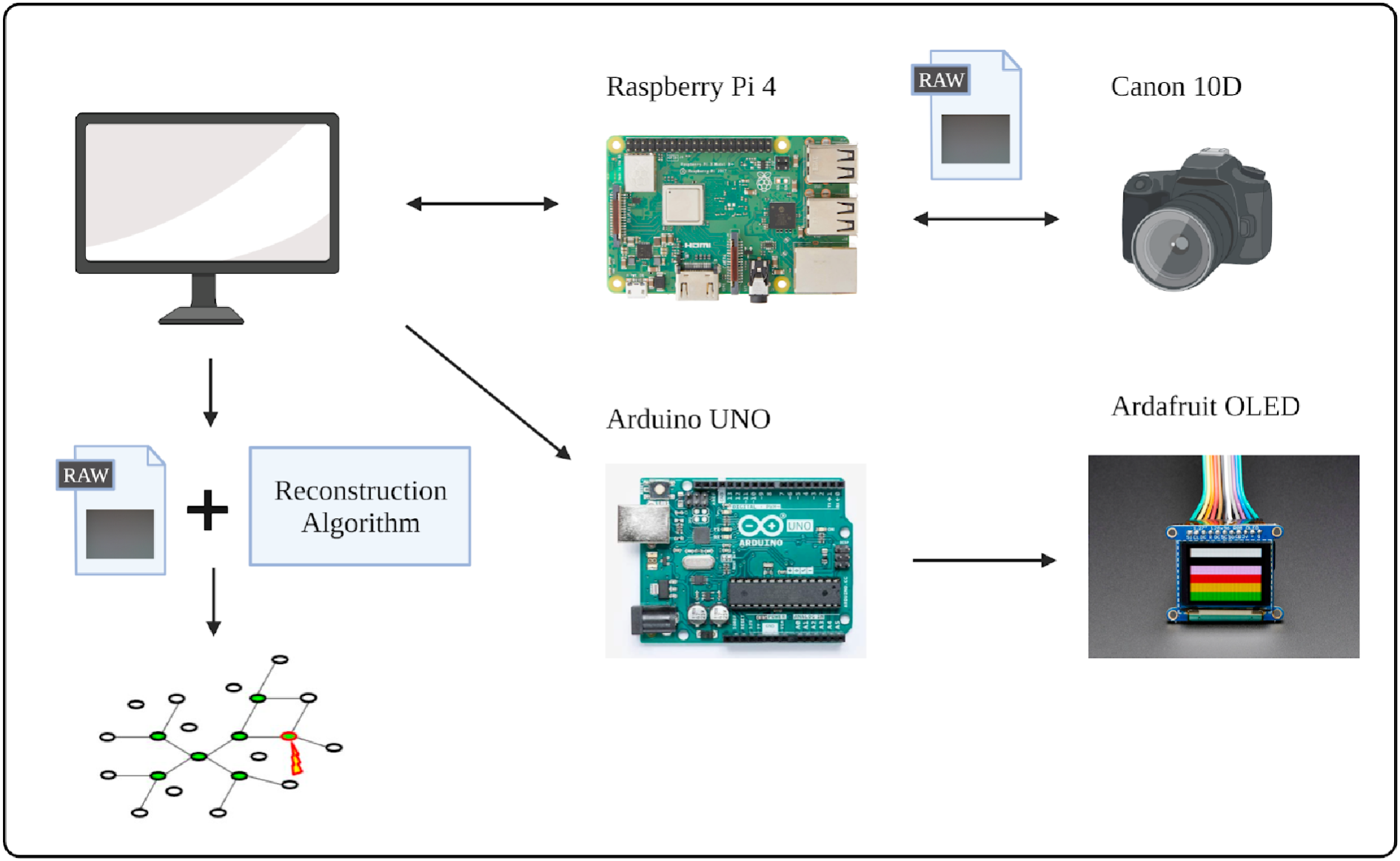
Integrated configuration for controlled illumination and image acquisition. Raspberry Pi and Arduino UNO are the board-level processors connected to a DSLR camera CMOS sensor and the OLED screen, respectively. Arrows indicate the directionality of signal transmissions. After triggering the capture function of the DSLR camera, the resulting image file is generated in the RAW format and then sent back to the PC for post processing and analysis.

### Software Architecture and Protocol Establishment

To provide synchronous control over the hardware setup, software architecture, and corresponding protocols were developed. As described in the hardware components section, the central controlling unit is the main User Interface (UI) application developed in the C# Windows Forms environment. The general functions of this application include: initializing serial port communication for the Arduino Uno, loading illumination settings into Arduino Uno, passing user-defined time-lapse imaging parameters to Raspberry Pi, and transferring files from Raspberry Pi to the PC. For the simplicity of future modifications and expansions, independent communication protocols were written between the two microprocessors and the main controlling unit, a PC in this case. The following sections will explain these parts in detail.

### Configuration of the Illumination Panel with the Arduino Uno

In this setup, digital strings that encode locational and wavelength information of the user-defined illumination pattern are streamed from the PC to Arduino using a USB serial connection. The USB port used for this data transmission is initialized once the UI application starts. Then, the AD2OLED.ino script, uploaded and contained in the Arduino memory, will constantly scan and decipher valid serial inputs. The encoding and deciphering algorithm uses a relatively straightforward structure that compiles the information into specific character sequences (**Figure 6**). Since the OLED panel used in the setup has a resolution of 128 x 96, all geometrical shapes can be decomposed into a series of rectangles with x and y coordinates and corresponding widths and heights represented by 3 digits. Once this information is successfully extracted from the string, the Arduino Uno makes function calls with these parameters to display the programmed pattern. With this serial communication process, the light source can be configured precisely with millisecond precision for optogenetics applications.

**Figure 6:**
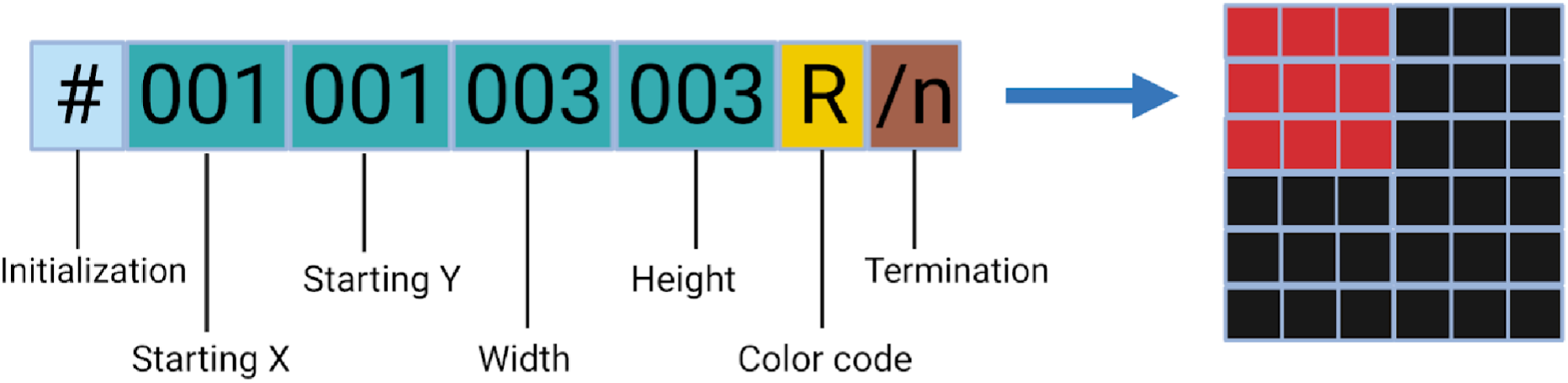
Data structure encoded for OLED display information.

### Camera Control and File Transferring using the Raspberry Pi

To decrease latency in data transmission and increase the stability of the system, a two-way communication system between the PC and Raspberry Pi is established through an ethernet connection. The oversight and encryption of this ethernet connection are done from the PC side using the PuTTY Secure Shell Protocol (SSH) platform. Besides the actual PuTTY client itself, two packages are needed for this application: (1) plink, a command line interface for remote execution of a python file on the Raspberry, and (2) pscp, SSH encrypted file transfer. The actual implementations of these two packages are done by specific event-handling scripts: PC2Pi_CC.py and PC2Pi_FT.py (CC stands for Camera Control and FT stands for File Transfer). Through the camera control script, time-lapsed imaging settings, such as the total number of shots and time interval in between shots, are parsed into the recipient script, ImageCapture.py, on the Raspberry Pi as input parameters. This recipient script adopted functions from the gphoto2 library to interface with the Canon 10D camera (gphoto2 supports digital access of many commercially available DSLR cameras). The final software architecture combined with all the protocols is shown in **Figure 7**. And the final main UI is shown in **Figure 8**. These files are available on GitHub https://github.com/BreakLiquid/Neural_Tracing.git.

**Figure 7:**
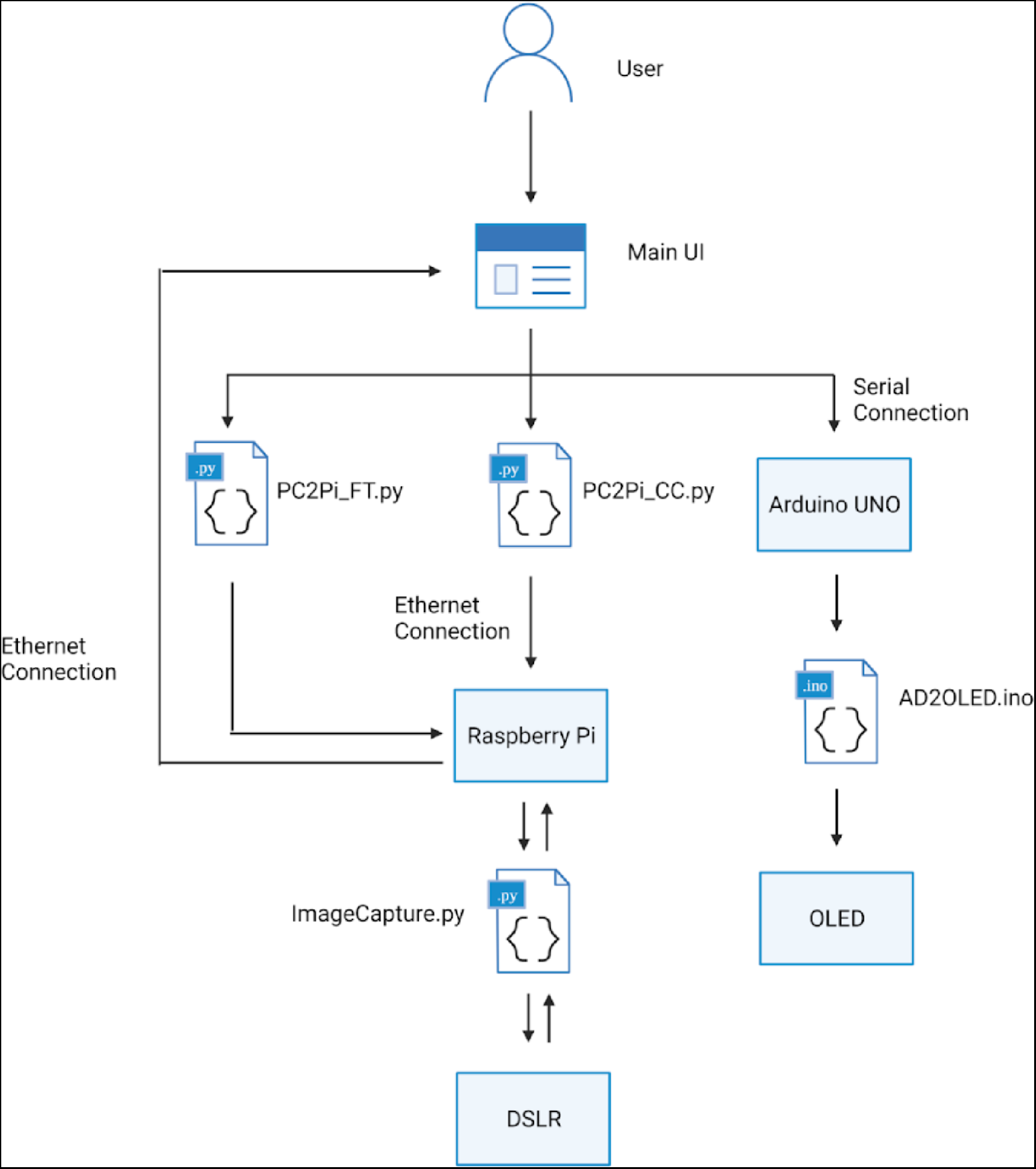
Overview of the software architecture. Functions like generating the desired illumination on the OLEO panel, capturing images with OSLR, and transferring image files from Raspberry Pi to PC are initialized at the main UI. For the OLEO control, the user inputs are converted into an information encoded string and sent to the Arduino UNO through the serial 1/0. A pre-loaded Arduino script will decipher the input string and output corresponding pattern alteration. For camera capture (CC) and file transfer (FT), an ethernet connection is used.

**Figure 8:**
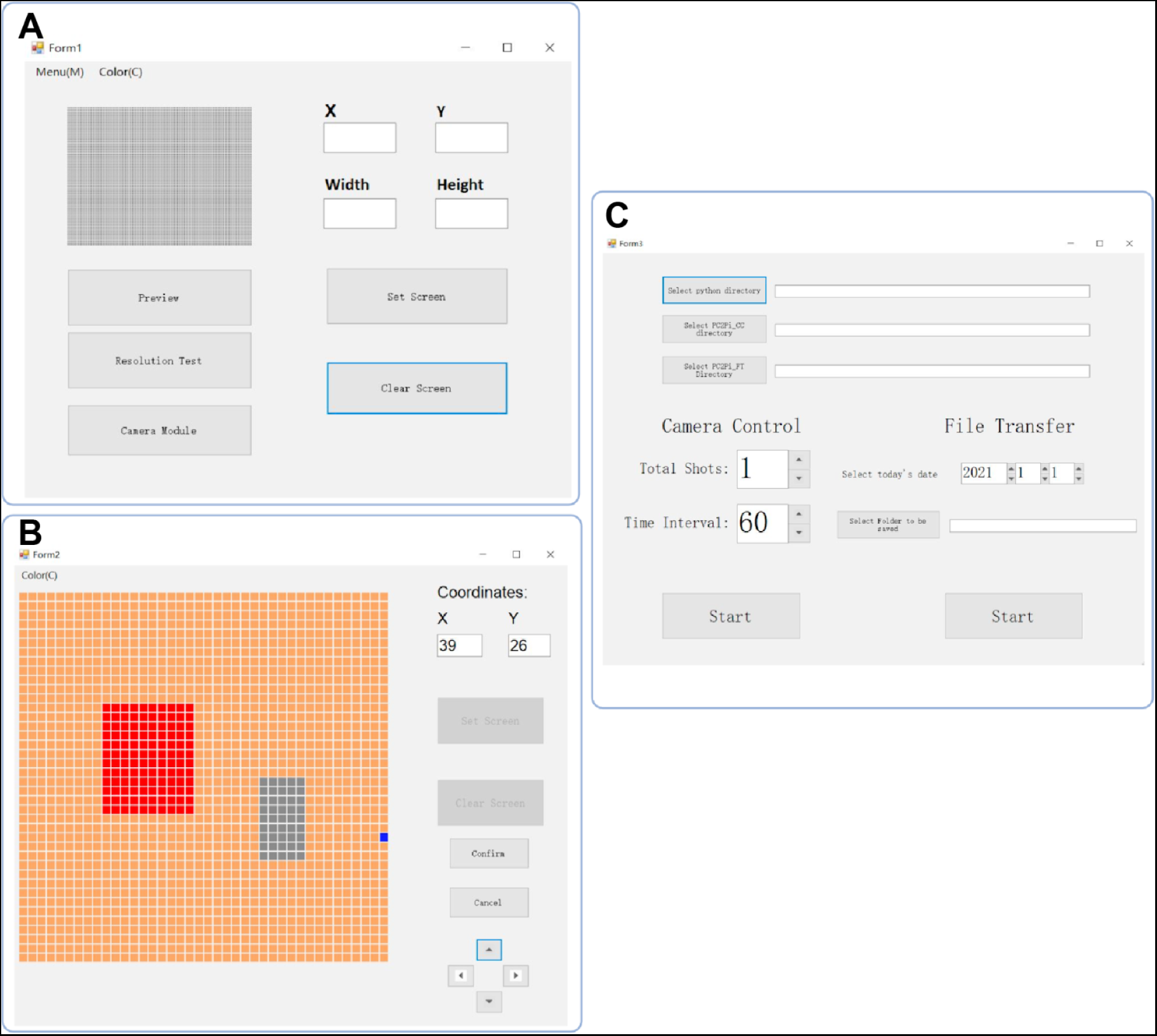
The user interface of the main application. **(A)** This is the initial form that will be shown once the application is opened. The pixelated map on the top-left represents the current display of the panel. Digital inputs on the right allow some basic configurations of the OLEO. This form also provides options such as resolution testing and clear screen. From this window, users can also open the other two forms. “Preview” button leads to Form2 and the “Camera Module” leads to Form3. **(B)** The canvas on the right is the map for pixel selection which can be done by a click and drag motion of the cursor. The current cursor location is highlighted in blue and its coordinates are displayed on the top-right corner. The red region represents the active pixel area. The grey region represents the area selected, but not yet active. Users can navigate through different sections of this panel using the arrow keys. **(C)** This form sets up the parameters for image capturing and transferring. Since all the image files saved on the Raspberry Pi are labeled by the time the image is taken, the transferring process requires the time information to locate the files.

To control both hardware devices synchronously, a software framework that connects the OLED panel and CMOS sensor with their corresponding microcontrollers, Arduino and Raspberry Pi, was constructed. With specified communication protocols, the end user can pass on commands such as triggering time-lapse imaging and changing displays with a user-friendly interface built on a PC platform. The image file transfer between the sensor module to the PC is also protected with a secure shell protocol.

## MATERIALS AND METHODS

The workflow of DNA plasmid construction for this project is shown in **Figure S2**. The DNA part assembly design for Golden Gate/uLoop assembly was done using a Google Sheet and with the open-source Multi-platform DNA editing software, A Plasmid Editor (APE) (51). After designing the high-level construct in the Google Sheet, low-level parts were designed on the sequence level using APE, such that they could be assembled using a GoldenGate-based DNA (52) assembly method uLoop (53) into the final construct. Low-level parts are produced and combined with uLoop (53) and then transformed into *E. coli* for sequence analysis and subsequent assembly into larger parts (see supplementary methods for details).

### Bacterial Transformation with Electroporation

For each assembled reaction, one vial of 30µL electrocompetent cell is thawed on ice and then mixed with 0.4µl of loop assembly plasmid using a 1mm electroporation cuvette. Immediately after electroporation, 100µL of SOC media is added into the cuvette, mixed, and then transferred into a 1.7 ml microcentrifuge tube. Electroporated cells are then incubated by shaking at 37 ℃ for 15min-1 hours. After the incubation, the mixture is spread on the Bluo-Gal (300ug/ml)+IPTG (1mM)-added LB antibiotic plate to allow colonies to grow overnight.

### Cre reporter assay in E18 Rat cortical neurons

E18 Rat Cortex was purchased from BrainBits/Transnetyx (Cat #SDECX). Neurons were disassociated using the Papain Dissociation System from Worthington Biochemical Corporation. Briefly, E18 Rat Cortex was incubated in DNAse+Papain mixture for 5-10 minutes at 37C with occasional gentle mixing by hand. Tissue was then triturated using a silanized 10ml glass pipette no more than 10 times. After letting undispersed pieces settle for 1 min, the supernatant was transferred into a new tube and centrifuged for 1 min at 200G. Cell pellets containing neurons were resuspended in NbActiv1 and then counted using a hemocytometer. Cells were then prepared for Nucleofection using a Lonza 4D-Nucleofector with the X unit. Cells were centrifuged for 1min at 200G and resuspended in P3 Primary Cell Nucleofector Solution 3 with Supplement 1, such that 250,000 cells/120ul was used for each electroporation condition. 1000ng of total DNA was used, which contained: 500ng of Cre reporter (pPK-338), 400ng or “filler DNA” expressing Gal4 (pPK-101) and 100ng of iCre plasmid (Addgene #116755). Nucleofection was performed using the EM-110 nucleofector program. Following nuceofection, cells were plated into a 24-well plate on poly-lysine coated coverslips in NbActiv1. The media was changed every 3 days. Neurons cells matured (10 days) they were fixed by adding 37% formaldehyde for 10min before mounting. Neurons were imaged with an Olympus BX53 fluorescent microscope using 488-GFP/568-RFP filters, a 10X UPlanFLN objective, and a Retiga 2000R camera.

### Hardware Configuration

To design a system that can be off the shelf, scalable, and to minimize cost, the following commercially available components were chosen for this project:

### OLED light panel

The illumination equipment used was the 1.27" OLED Breakout Board with 16-bit color control from Adafruit. This active display panel comprises 128 x 96 RGB pixels with a 250µm x 250µm pixel size.

### CMOS sensor

The CMOS sensor used was from a Canon EOS 10D, 6.3 megapixel DSLR, with an effective imaging area of 22.7mm x 15.1mm, and a pixel pitch of 7.38µm.

### Hardware control/software

For the convenience of integrating post-processing with computational algorithms in the future, all the electronics are communicated through a Windows Form-based application written for a PC system. We designed the system such that users can pass in defined parameters into microprocessors used to control the illumination pattern on the OLED and to forward commands to the DSLR camera CMOS sensor for image capturing, RAW image file extracting, and file transferring from DSLR to PC. The microprocessors used to achieve these features are an Arduino UNO board and a Raspberry Pi 4 for the OLED and DSLR respectively (**Figure 5 and Figure 7**).

## DISCUSSION

We have constructed plasmids for controlling RV spreading using red and far-red light with TN7 sites for easy insertion into BACs. Using the mammalianized BV vector system (54) (55), it is possible to package these into viral vectors that can control RVΔGL versions of rabies, which is a considerable challenge with many other vectors given the size of the L gene alone. The BV genetic cargo capacity allows incorporating these genes as well as Cre reporters, insulators, optogenetic tools, and more. Importantly, the positive control constructs containing the Cre reporter and *hSyn-driven* L gene packaged into a mammalianized BV vector would be useful for mono-synaptic tracing using RVΔGL-Cre. Previously, RVΔGL-Cre could only infect a primary neuron because the L gene is too large (6.3kb, not including the promoter) to package into AAV, which is typically used to control the monosynaptic spreading of RV. The constructs we generated in this work are ready to insert into any system containing a TN7 insertion site, such as the Bac-to-BAC or MultiBacMam baculovirus systems.

To test and optimize the BV tracing system, neuronal cultures can be infected with BV following a low titer of RVΔGL-Cre. Illumination with red and far-red light from the OLED panel or using LEDs from Kyriakakis et al. (25,56) can test and optimize ratios, titers, leakiness, *etc.* Control experiments can be set up with different varying parameters such as the *L* gene *vs* destabilized *L* gene and different pulse lengths for the red light activation. Since the PhyB system can be shut off with far-red light, the *L* gene can be minimally expressed, minimizing cellular toxicity. If TVA-expressing mice are used as a source of organotypic brain slices, RVΔGL-Cre pseudotyped with RABVG could be injected *in vivo* in specific locations and infect specific cell types before slice preparation. This would allow for selecting a precise location *in vivo* for the starter cells and then multiple slices from the same brain could be traced using optogenetics.

### Using Golden-Gate assembly to build complex gene circuits

The uLoop system is an excellent system for building the complex circuits used in this work. This is particularly the case when needing to build different versions of a circuit. For example, when we built the *destabilized L gene* version of our ∼30kb construct, we first modified a smaller plasmid containing the L gene. This additional part was then used to re-assemble the rest of the construct with existing parts from the original design. Since uLoop is recursive, it facilitates building large genetic circuits as in this work would be difficult or impossible to build using other Golden Gate assembly methods.

### Optoelectronic for facilitating scalable and automated viral tracing

To facilitate control of the optogenetic switch and rabies tracing, our lensless imaging, and stimulation design would allow for inexpensive and scalable tracing experiments. The stimulation and imaging parts are composed of commercially available OLED panels and DSLR CMOS chips. These devices will only become cheaper, will have higher resolution over time, and can even incorporate microlenses and other novel optics that would further increase the capability of this approach (57–59). While a lensless imaging approach to imaging cells infected with BV and Rabies may be limited in resolution and sensitivity compared to other imaging methods, it enables the ability to image and stimulate a large FOV in real-time with a much smaller footprint than a microscope, similar to Pollmann *et al.*(*60*). After optogenetic/tracing, samples can be imaged using other modalities to enable deeper analysis. For example, samples could be fixed and immunostained to obtain a higher resolution with cell-type-specific markers. Samples could also be imaged using single-cell spatial imaging approaches that use the Cre reporter RNA and the RV RNA as labels. Since the samples can be post-processed and analyzed in such ways, the trade-offs of using a lensless imaging system for tracing are minimal. When applied to viral vector tracing, this setup will allow users to modify the illumination display for spatially specific activation of the optogenetics switches (controlling viral spreading, any gene of interest or neural activity) in real-time without the need of taking out the neuron samples in and out of their incubation environment or a microscope.

### Challenges to combining parts required for automated neural tracing

To achieve automated optically guided synaptic tracing, three core aspects of this work must be combined. First, high titer BV containing the tracing tools needs to be produced and tested in neurons to optimize control of RVΔGL. Second, the illumination system would have to be optimized for tracing. Third, the imaging component needs to be combined with the illumination system and cells/tissue to optimize the computational approaches to imaging. Next, the concept of spatially controlling the spread of RV using OLED stimulation can be tested. Since PhyB optogenetic control of cellular activity has been done on a subcellular level in previous studies (28,29), single-cell control of transsynaptic spreading should be feasible. To achieve this using an LED array would require calibrating the parameters associated with the neural tracing applications such as the titer of BV cells, the intensity of the red illumination, and suppression with wide-field far-red illumination. For post-imaging analysis, there is still a need to develop an image reconstruction algorithm that fits well with the fluorescent signal of the neuron sample. Since the fluorescent signal used to target the BV-infected starter neuron is confined to the nucleus and optogenetic stimulation is only required in the nucleus, this level of precision should be achievable. One promising method for locating BV-infected neurons is to use a Point-Spread-Function (PSF) estimation on the light source and deconvolve the raw image into useful spatial information (61) (62). This method has been applied in numerous lensless imaging applications achieving robust reconstruction results. This conceptual design could be adapted using higher resolution LEDs with larger panels and combining multiple CMOS sensors to trace larger brain slices or multiple brain slices on a single setup. Further, many of these devices can easily fit into a standard cell culture incubator, allowing mapping sections of entire brains or multiple brains.

### Applications of optogenetics and other Genetically-Encoded and Physically-Actuated (GEPA) tools for controlling viral tracing tools

Our optogenetically guided approach to tracing could also be used for other viral tracing tools, where genes are identified as necessary and sufficient to rescue the transsynaptic spreading function when expressed *in trans.* For example, light could modulate the expression of the Herpes Simplex Virus strain H129 thymidine kinase/TK (63), RABV-G in the CVS-N2c strain (64), or RABV-L (65). Further, the optogenetic tools and reporters can be expressed using more specific cell-type specific promoters, restricting the tracing to neuronal subtypes (66). Using a high-capacity vector that can also trace neuronal connections, such as HSV, it may be possible to package the entire optogenetic switch into the vector allowing for transsynaptic tracing using a single virus. Similarly, other modalities to control genes through external stimulation (Genetically-Encoded and Physically-Actuated (GEPA) tools), such as sonically/mechanically (67,68), or magnetically (69), may be used as guides to trace circuits. These tools would allow for the delivery of genes in a spatially confined way that is cell-type specific and into specific circuits. This capability could facilitate controlled polysynaptic tracing *in vivo* or for stimulation-guided gene delivery, which could have therapeutic potential to deliver genes into specific circuits in the body/brain from the periphery.

## Acknowledgments

This work was supported by the Kavli Institute for Brain & Mind and NIH/ NIDCD grant 1R21DC018237-01. We want to thank Dr Fernan Federici and Dr. Jim Haseloff for providing the uLoop plasmid system. We also want to thank Dr. Imre Berger for his helpful discussions about the MultiBACMam system.

## SUPPLEMENTARY MATERIALS

### DNA Fabrication with uLoop Assembly

Construction of plasmids containing the expression cassettes for baculovirus vectors was done through uLoop assembly detailed in Pollak *et. al.* (*53*). This Type IIS restriction enzyme-based DNA assembly method can generate large constructs of DNA plasmids (∼30kb in this work) by recursive DNA assembly (**Figure S3 and S4**). In the uLoop method, the Type IIS restriction enzymes used are BsaI and SapI. Type IIS restriction enzymes like Bsa1 and Sap1 make DNA cuts from their recognition sites, allowing cutting off the DNA while removing the Type IIS site and generating customized hangover sites, essential properties for Golden Gate-based DNA assembly methods (**Figure S3**). In uLoop, BsaI and Sap1 restriction sites are arranged in a manner, which is reversely oriented in Odd (L1, L3, *etc.*) and Even (L2, L4, *etc.*) levels (53). To produce level 0 parts (L0), the basic gene elements are “domesticated” and then assembled into the Odd receivers plasmids. This domestication process involves removing BsaI and SapI sites from the wild-type gene sequences to prevent cutting the parts during the assembly process. For large DNA elements such as the *L* gene, multiple mutations were required to achieve complete domestication. Thus, the C-terminal part of the *L* gene was synthesized (IDT gBlock) using different codons. In the next step of uLoop, four L1 plasmids are assembled into an Even receiver plasmid to make a level 2, or L2 plasmid **(Figure S4)** (53). During this process, the SapI enzyme cleaves the parts out of L1 plasmids and removes the *LacZ* marker in the Even receiver. Then, the four compatible L1 parts/overhangs ligate to each other and form an L2 plasmid containing desired parts in a predetermined orientation (**Figures S3 and S4**). This process is repeated for the assembly of L3 plasmids, using the BsaI restriction enzyme instead of Sap1. L4 plasmids can then be produced using the SapI enzyme again (**Figure S4**). The detailed reagents and protocols used for the assembly were adopted from the Pollak *et al.* study (*53*).

---

**Figure S1:**
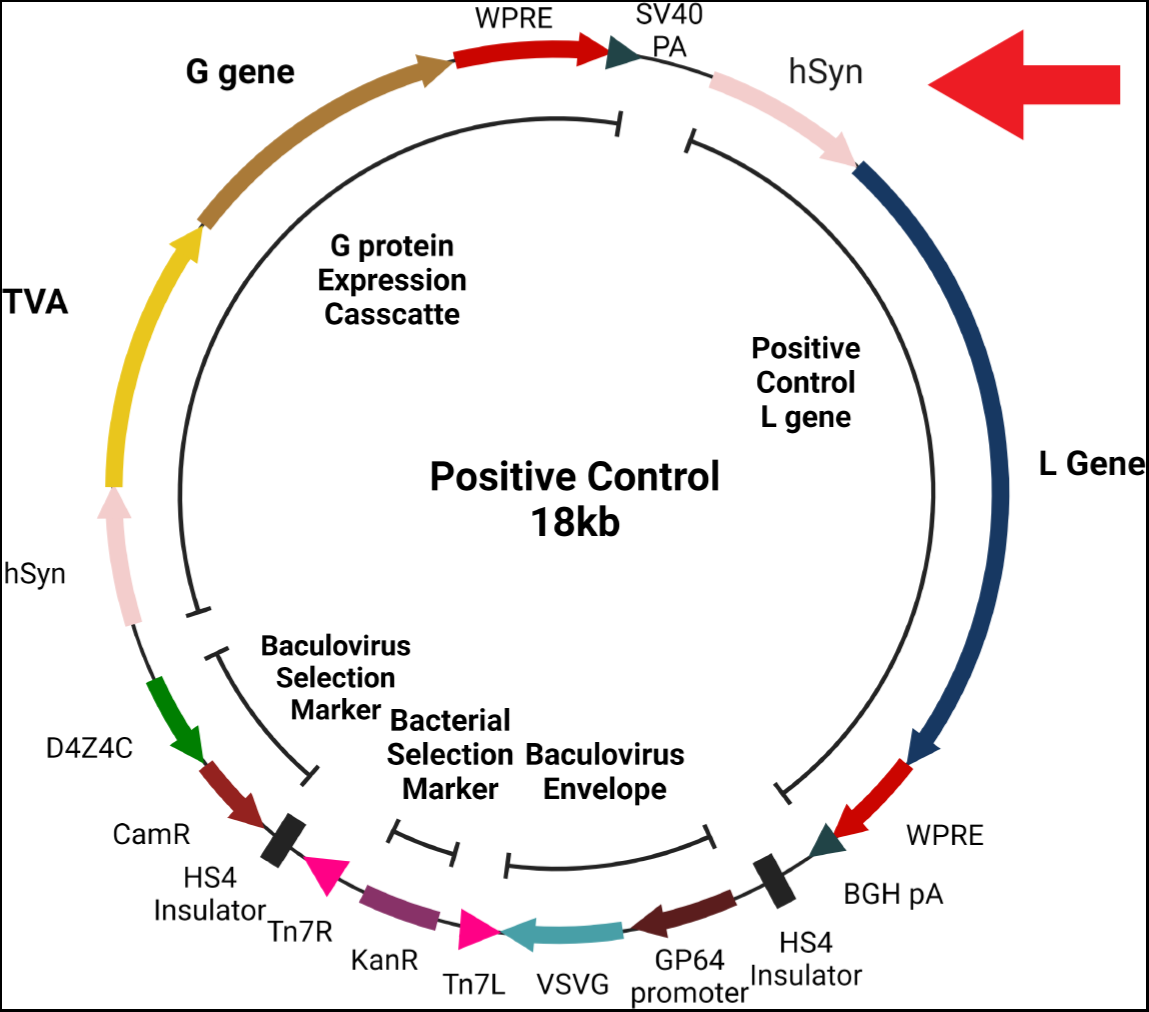
Positive control plasmid for aGL RV tracing. The positive control plasmid contains the L gene under the control of the constitutive hSyn promoter (Red Arrow) instead of a light inducible promoter. (CamR = Chloramphenicol Resistance, D4Z4C = insulator, hSyn = Synapsin promoter, WPRE = Woodchuck hepatitis virus Post-transcriptional Response Element, BGH PA= Bovine Growth Hormone PolyA tail, SV40 PA= Simian Vacuolating virus 40 PolyA tail, KanR = Kanamycin Resistance)

**Figure S2:**
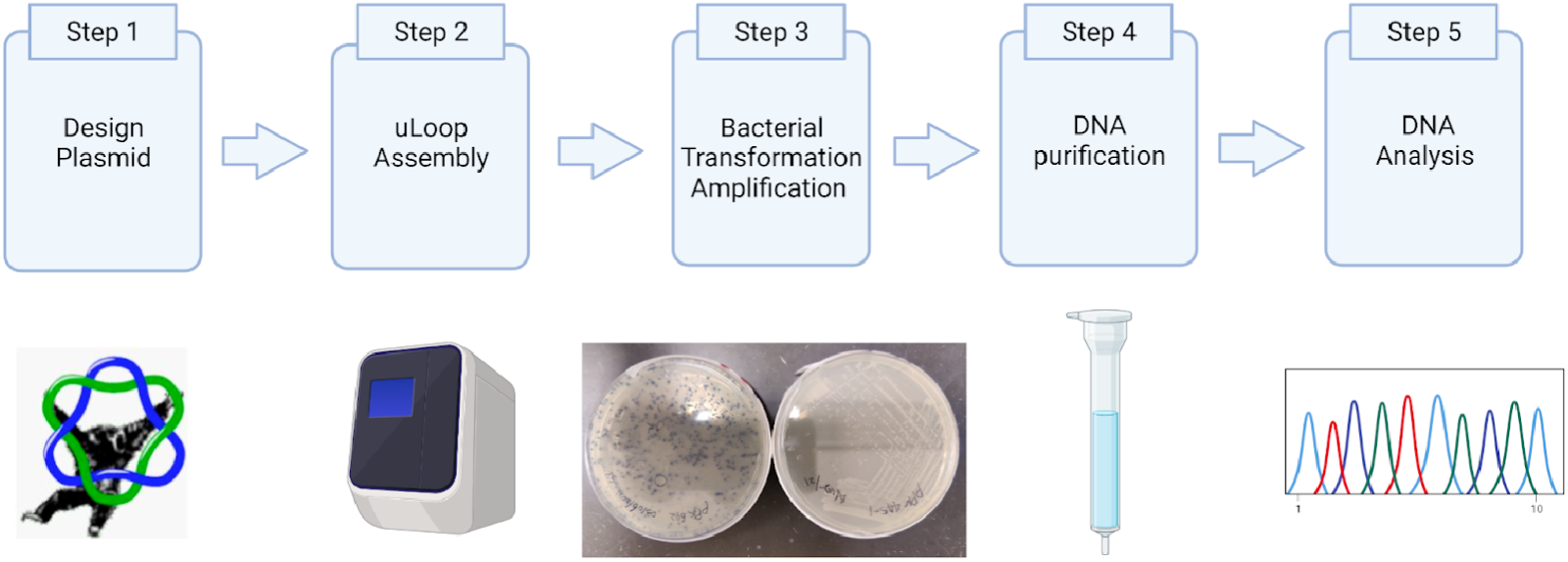
Overall workflow of the plasmid construction.

**Figure S3:**
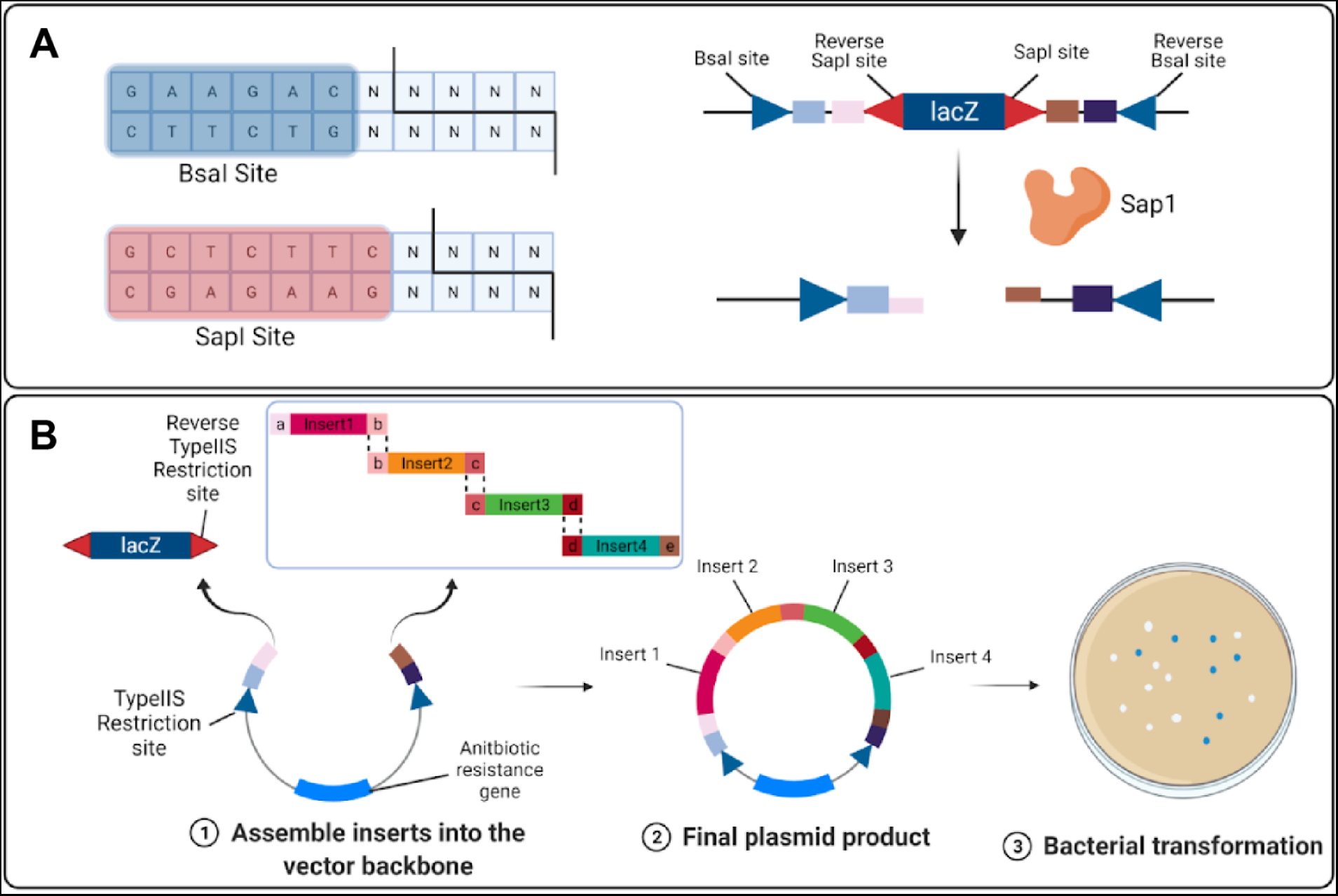
uloop assembly with Lacz implementation for blue-white selection. **(A)** Bsal and Sapl are the two Type IIS restriction enzymes used for uloop assembly. Their corresponding recognition sites are labeled in blue for Bsal and red for Sapl. The backbone of the uloop assembly uses the property that Type IIS enzymes cut outside the recognition sequence to generate customized single-stranded overhangs for position-specific DNA part assembly. The inverted placement of these restriction sites allows the enzyme to cut out the lacZ gene from the backbone plasmid and leave overhangs that match the DNA to be inserted. **(B)** During the uloop assembly process, four DNA elements in the same level (Ln) are cut with the corresponding enzyme to make four linear DNA fragments. Since the Type IIS enzymes leave the inserts with matching overhangs that reflect their relative positions in the final assembled plasmid, these inserts only ligate in this specific alignment: insert1-insert2-insert3-insert4. The restriction enzyme also cuts out the lacZ gene from the next level (Ln+1) backbone and leaves overhangs for the final insert combination. The successfully assembled plasmid product includes the correct antibiotic selection gene for that level while excluding the lacZ gene. LB agar plates containing the antibiotic along with BluoGal+lsopropyl f3-D-1-thiogalactopyranoside (IPTG) blue-white screening, which differentiate the colonies with wanted plasmid result (white) from the colonies with the unwanted backbone (blue), are used to select for assembled parts.

**Figure S4:**
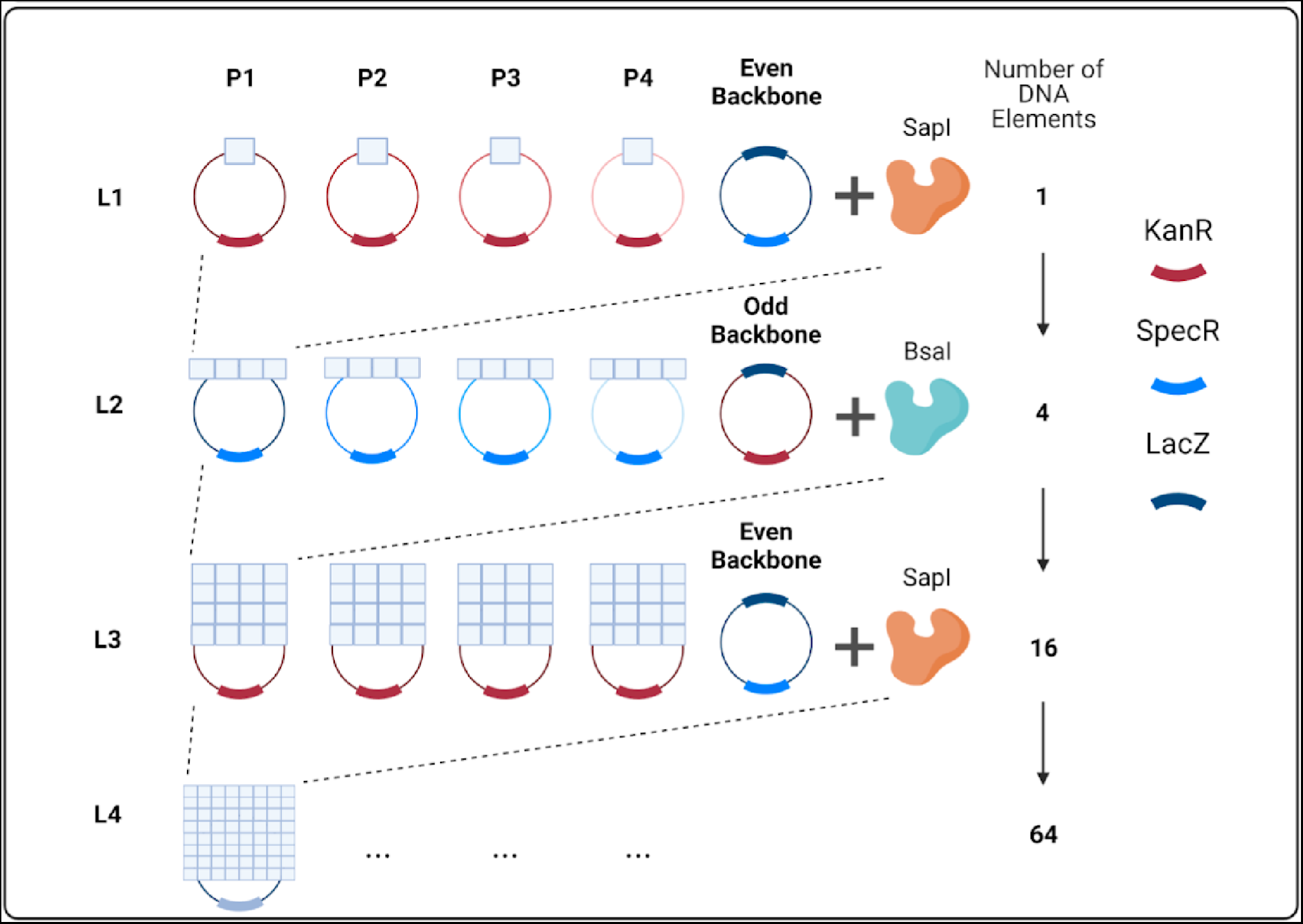
Recursive DNA assembly using uloop. The overall assembly process follows this trend where Sapl is used for making even level parts (L2 and L4) and Bsal is used for making odd level (L3) parts. The antibiotics selection markers are also separated in a similar way with KanR on the odd level plasmids and SpecR on the even level plasmids. The level of the plasmid also reflects the number of DNA elements in the plasmid.

**Figure S5:**
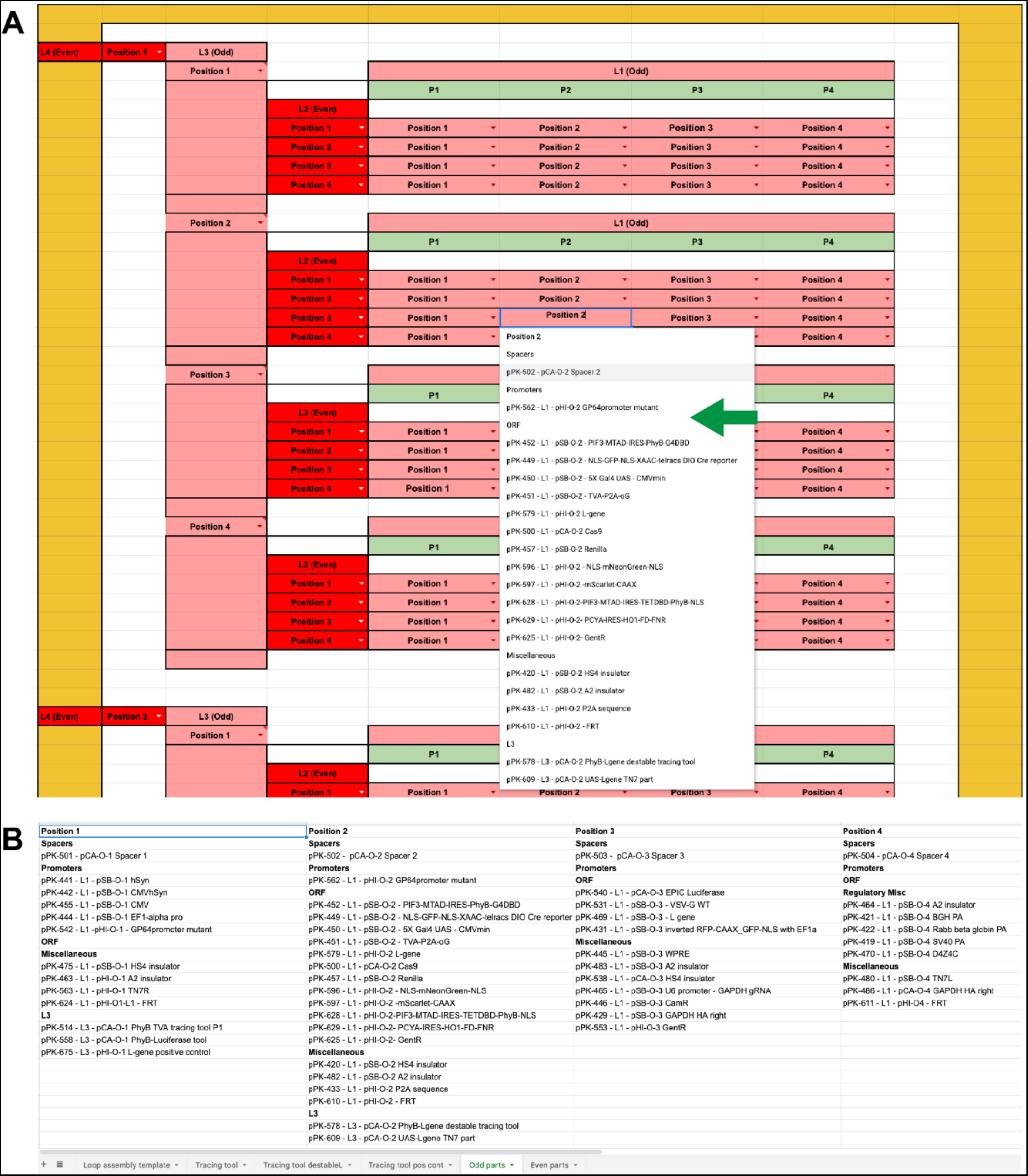
A spreadsheet used for planning and tracking uloop constructions. **(A)** The empty template used for planning uloop constructs_The (Green Arrow) points to the drop down menu that appears when a position is selected. **(B)** The tab containing parts is linked to the template for creating a customizable drop down menu.

**Figure S6:**
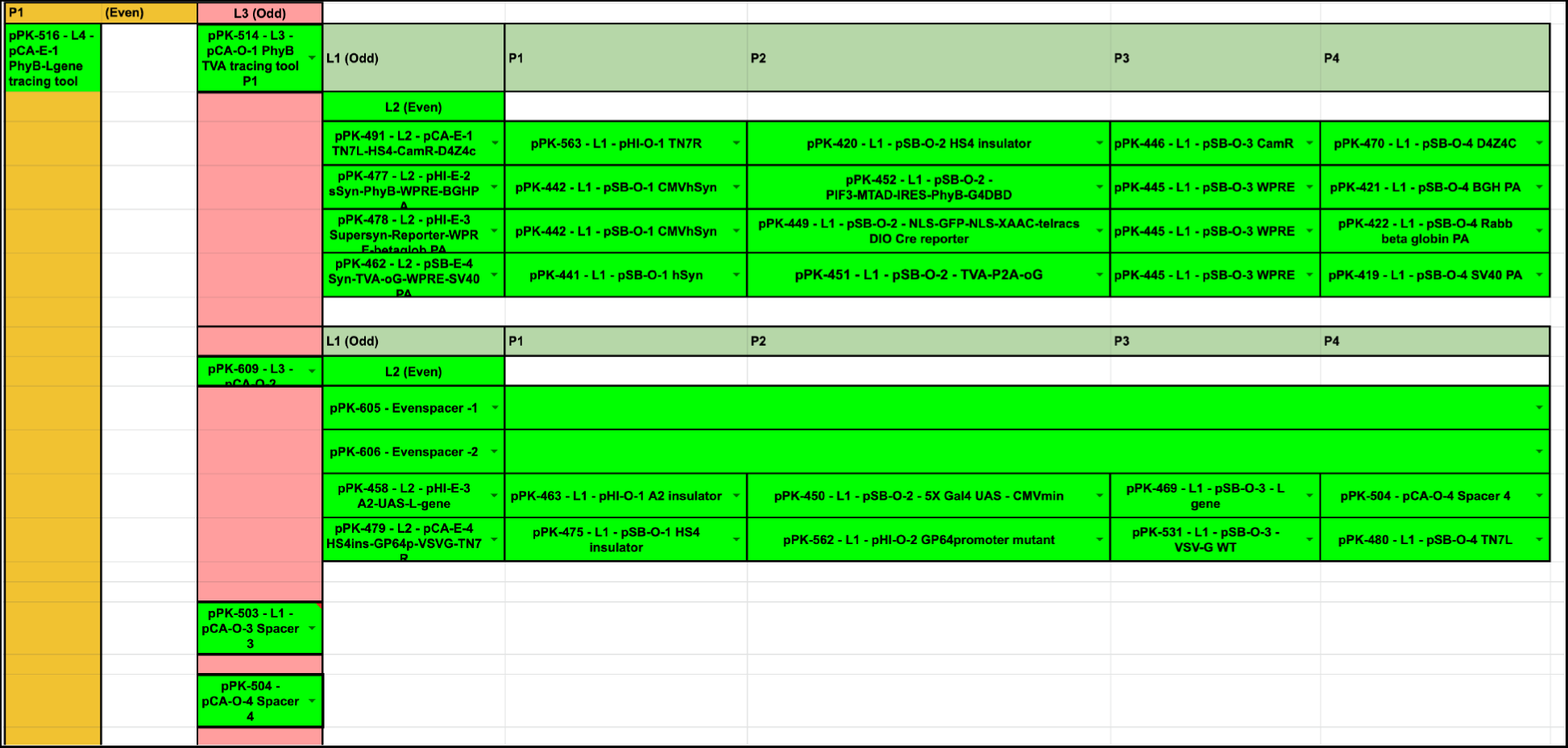
An example construction mapped out using the uloop assembly template. **(A)** The level 4 (L4) part pPK-516 is constructed of two L3 parts (pPK-514 and pPK-609) and two spacer parts (pPK-503 and pPK-504). The level 3 part (L3) pPK-609 is constructed from spacer parts (pPK-605 and pPK-606) and two level 2 (L2) parts (pKP-458 and pPK479). Using the drop down menus starting with level 1 parts, existing parts can be chosen to design level 2 parts. Coloring the boxes can indicate progress in the construction (we used green to indicate completed parts and red to indicate incomplete parts).

## Notes

https://github.com/BreakLiquid/Neural_Tracing.git

### Competing Interest Statement

The authors have declared no competing interest.

### Summary of Updates

A missing citation was added, and an acknowledgments section was added.

